# The human innate immune protein calprotectin elicits a multi-metal starvation response in *Pseudomonas aeruginosa*

**DOI:** 10.1101/2021.05.04.442697

**Authors:** Cassandra E. Nelson, Weilang Huang, Emily M. Zygiel, Elizabeth M. Nolan, Maureen A. Kane, Amanda G. Oglesby

## Abstract

To combat infections, the mammalian host limits availability of essential transition metals such as iron (Fe), zinc (Zn), and manganese (Mn) in a strategy termed “nutritional immunity”. The innate immune protein calprotectin (CP) contributes to nutritional immunity by sequestering these metals to exert antimicrobial activity against a broad range of microbial pathogens. One such pathogen is *Pseudomonas aeruginosa*, which causes opportunistic infections in vulnerable populations including individuals with cystic fibrosis. CP was previously shown to withhold Fe(II) and Zn(II) from *P. aeruginosa* and induce Fe- and Zn-starvation responses in this pathogen. In this work, we performed quantitative, label-free proteomics to further elucidate how CP impacts metal homeostasis pathways in *P. aeruginosa*. We report that CP induces an incomplete Fe-starvation response, as many Fe-containing proteins that are repressed by Fe limitation are not affected by CP treatment. The Zn-starvation response elicited by CP seems to be more complete than the Fe-starvation response and includes increases in Zn transporters and Zn-independent proteins. CP also induces the expression of membrane-modifying proteins, and metal-depletion studies indicate this response results from the sequestration of multiple metals. Moreover, the increased expression of membrane-modifying enzymes upon CP treatment correlates with increased resistance to polymyxin B. Thus, response of *P. aeruginosa* to CP treatment includes both single and multi-metal starvation responses and includes many factors related to virulence potential, broadening our understanding of this pathogen’s interaction with the host.

**Importance:** Transition metals are critical for growth and infection by all pathogens, and the innate immune system withholds these metals from pathogens to limit their growth in a strategy termed “nutritional immunity”. While multi-metal depletion by the host is appreciated, the majority of metal depletion studies have focused on individual metald. Here we use the innate immune protein calprotectin (CP), which complexes with several metals including iron (Fe), zinc (Zn), and manganese (Mn), and the opportunistic pathogen *Pseudomonas aeruginosa* to investigate multi-metal starvation. Using an unbiased label-free proteomics response, we demonstrate that multi-metal withholding by CP induces a regulatory response that is not merely additive of individual metal starvation responses, including the induction of Lipid A modification enzymes.

## Introduction

Transition metals are essential for all life, and invading microbial pathogens must acquire these nutrients to grow in the host and cause infection. The host innate immune system limits growth of microbial pathogens by withholding essential transition metals through a strategy termed “nutritional immunity” (1, 2). Metal-sequestering innate immune proteins are important components of this host response. Nutritional immunity originally focused on the competition for ferric iron [Fe(III)] wherein host proteins such as lactoferrin sequester Fe(III) and bacterial siderophores scavenge Fe(III) from these host proteins and deliver it to bacteria via siderophore receptors (1). Later, with the discoveries of additional metal-sequestering host proteins and microbial metal-uptake systems, the model for nutritional immunity expanded to include other nutrient metals such as Mn(II) and Zn(II) (2, 3). The S100 protein calprotectin (CP, S100A8/S100A9 oligomer, MRP8/MRP14 oligomer) plays a central role in nutritional immunity because it sequesters multiple divalent metal ions including manganese [Mn(II)], iron [Fe(II)], nickel [Ni(II)], and zinc [Zn(II)] [reviewed by (4)]. CP exerts antimicrobial activity against a broad range of bacterial and fungal pathogens, and this activity is generally attributed to its metal-withholding ability (5–10).

Because the host deploys multiple metal-withholding proteins that coordinate various metal ions (e.g. CP, lactoferrin, siderocalin, S100A12) at infection sites, bacterial pathogens must respond to the concerted limitation of multiple metal nutrients in order to cause infection (11). Moreover, bacterial pathogens exhibit varying nutritional requirements for transition metals – for instance, some species have a high Fe requirement whereas others require substantially more Mn (12, 13) – and are therefore likely to show species- and strain-specific responses to host multi-metal withholding. Indeed, a recent study of Gram-negative and Gram-positive bacterial pathogens revealed that the effects of CP on the uptake of nutrient metals depends on the organism and the culture media (10). Despite the current appreciation for multi-metal sequestration by the host, the microbial response to this stress has received relatively less attention than single-metal limitation (11). Because CP sequesters multiple metals, it provides a physiologically important and useful tool for studying the response of microbial pathogens to multi-metal withholding by the host.

*Pseudomonas aeruginosa* is a Gram-negative opportunistic pathogen that causes infections in vulnerable populations, including individuals with cystic fibrosis (CF). *P. aeruginosa* has a high Fe requirement and uses multiple Fe-uptake systems to acquire Fe from the host environment including siderophore-mediated Fe(III)-uptake systems, an Fe(II)-acquisition system, and heme-uptake systems [reviewed by (14)]. Due to the propensity of excess Fe to cause oxidative damage, a complex and hierarchical Fe-regulatory system regulates the uptake of Fe in response to cellular Fe concentrations [reviewed by (15)]. This function is mediated by the Fe-binding transcriptional regulator Fur, which in its holo form binds to the promoters of genes encoding for Fe-uptake systems, thereby blocking their transcription and Fe uptake (16). As Fe levels decrease, Fe dissociates from Fur and apo Fur loses affinity for these promoters, allowing transcription of Fur-repressed genes. In addition to Fe-uptake genes, Fur also represses the PrrF small-regulatory RNAs (sRNAs), which post-transcriptionally repress the expression of nonessential Fe-containing proteins and Fe-storage proteins (17, 18). This so-called Fe-sparing response reduces the Fe requirement of the cell and is required for *P. aeruginosa* pathogenesis (19–21).

The mechanisms used by *P. aeruginosa* to maintain homeostasis of other transition metals are less well characterized, but several key aspects of select uptake and regulatory systems have been identified. The Zn(II)-homeostasis system in *P. aeruginosa* includes Zur, a Fur homolog, that represses the transcription of Zn(II)-uptake systems (22). *P. aeruginosa* has several characterized Zn(II)-uptake systems including the Znu permease, the pseudopaline-metallophore-mediated Cnt (also called Zrm) system, and the HmtA P-type ATPase, as well as putative ABC permease gene clusters (PA2911-PA2914, PA4063-PA4066) (23–26). Repression of Zn(II)-containing proteins has been shown under Zn(II)-limiting conditions, indicating a Zn(II)-sparing response (23, 26). When Zn(II) is in excess, transporters such as the resistance-nodulation-division (RND) pump CzcCBA (27), the cation diffusion facilitator transporters CzcD and ZitB (28), and the P-type ATPase transporter ZntA (29) efflux Zn(II) to prevent toxicity. In terms of Mn homeostasis, *P aeruginosa* encodes two predicted Mn(II) transporters (MntH1, MntH2) (30) and requires Mn for several Mn-dependent proteins, including the ureohydrolases (GbuA, GpuA) (31) and the superoxide dismutase SodM (32). While aspects of *P. aeruginosa* Mn regulation have been described (30, 33), it remains unclear how this pathogen maintains Mn homeostasis in the context of the host metal-withholding innate immune response. The Cu-homeostasis system has also been characterized in *P. aeruginosa* and is comprised of Cu importers, Cu chaperones, and Cu efflux pumps to maintain Cu homeostasis (34).

CP is found in high concentrations in the lungs of CF patients, which are commonly infected by *P. aeruginosa* (35–37). Moreover, CP has been shown to reduce the antimicrobial activity of *P. aeruginosa* toward another CF pathogen, *Staphylococcus aureus*, which was attributed to its ability to inhibit the production of toxic secondary metabolites by *P. aeruginosa*, including phenazines and 2-alkyl-4(1*H*)-quinolones (AQs) (38). We previously reported that CP withholds Fe from *P. aeruginosa* and thereby causes an Fe-starvation response through targeted analyses of known Fe-acquisition and -regulatory systems (10). Our study also demonstrated a change in virulence factor production in response to CP with the downregulation of the phenazines, which we determined to be a consequence of Fe limitation (10). Moreover, a recent investigation demonstrated that CP withholds Zn(II) from *P. aeruginosa* and induces a Zn-starvation response (39). This study also reported decreased Zn(II) protease activity caused by CP, indicating attenuated virulence potential. Because previous studies on the impact of CP on *P. aeruginosa* have focused on a single metal, we sought to determine the response of *P. aeruginosa* to multiple-metal withholding by CP.

In this work, we investigated how CP-dependent metal depletion affects *P. aeruginosa* physiology and virulence capacity using quantitative, label-free proteomics. In agreement with prior studies (10, 39), our analyses show that CP causes both Fe- and Zn-starvation responses in *P. aeruginosa*. Moreover, we identified increases in the expression of Zn(II)-containing proteases and membrane-modifying enzymes in response to CP. To decipher whether the observed changes in protein expression resulted from single-metal or multi-metal sequestration by CP, we used our proteomics workflow to evaluate the consequences of Fe, Mn, and Zn limitation on the *P. aeruginosa* proteome. This effort revealed that CP elicits an expected Zn-starvation response and what appears to be an incomplete Fe-starvation response. Moreover, CP treatment leads to the induction of membrane-modifying enzymes, likely as a consequence of multi-metal sequestration. This response correlated with increased resistance to polymyxin B, suggesting that host metal sequestration may induce *P. aeruginosa* resistance to cationic antimicrobial peptides (CAMPs). Together, this work shows that the response of *P. aeruginosa* to multi-metal withholding by CP is distinct from that of single-metal withholding and suggests a complex microbial response to this innate immune protein.

## Results

### CP causes Fe- and Zn-starvation responses and alters virulence protein production in *P. aeruginosa*

To determine the global response of *P. aeruginosa* to CP, we evaluated the impact of 10 μM CP treatment on the PA14 proteome. PA14 was grown under conditions that were previously used to study the consequences of CP treatment on *P. aeruginosa* and other pathogens (10, 40). Specifically, PA14 was cultured under aerobic conditions in a metal-replete chemically-defined medium (CDM) containing 2 mM Ca, 5 μM Fe, 0.3 μM Mn, 6 μM Zn, 0.1 μM Ni, and 0.1 μM Cu, in the presence or absence of 10 μM CP. These metal concentrations were selected to be within the reported range of metal levels in the sputa of CF patient samples (41–43), and the concentration of CP is within the range of CP concentrations identified at infection sites (44). We selected CDM for these studies because it allows for control over the metal concentrations and sources (e.g., non-heme vs heme Fe), and it is amino acid rich similar to what is found in CF sputum (45). Prior whole-cell metal analyses in metal-replete CDM revealed that treatment of PA14 with CP results in a significant decrease in cell-associated Fe and negligible change in cell-associated Mn, Ni, Cu, and Zn (46). PA14 was grown for 8 hours, which afforded growth to early stationary phase (10). Cells were harvested, and quantitative label-free proteomics was performed using nano ultra-performance liquid chromatography coupled to high-resolution tandem mass spectrometry as previously described (47–49). Protein levels that were significantly (p<0.05, n=5) changed at least 2-fold (equivalent to 1 log_2_-fold change or LFC) were analyzed further.

CP treatment caused a significant increase in the abundance of 93 proteins and a significant decrease in the abundance of 72 proteins (**Fig 1A**, **Table S1**). To identify biological connections within the upregulated and downregulated proteins, we performed network analyses using the STRING database (50). The networks created for proteins upregulated (**Fig 1B, Table S1**) or downregulated (**Fig 1C, Table S1**) upon CP treatment showed significantly more interactions than if they were a random collection of genes from the genome, indicating a biological relationship between the proteins in each analysis (**Table S2**). In agreement with recent studies showing that CP induces an Fe-starvation response in *P. aeruginosa* (10, 46), proteins involved in pyoverdine-mediated Fe(III) uptake and heme-dependent Fe acquisition were upregulated in response to CP, whereas proteins involved in phenazine biosynthesis (PhzE, PhzD, PhzC, PhzM, PhzH) and secretion (MexH, MexI) were downregulated (**Fig S1**). CP treatment also resulted in an apparent Fe-sparing response, indicated by the downregulation of nonessential Fe-containing proteins. For example, nonessential Fe- and heme-containing proteins such as LeuC and KatA were downregulated upon CP treatment, whereas the Fe-independent paralog of fumarase (FumC1) and the Mn-dependent superoxide dismutase (SodM) were upregulated in response to CP (**Fig S1**). However, CP treatment did not reduce levels of several Fe-containing tricarboxylic acid (TCA) cycle proteins, which were previously shown to be repressed by the PrrF sRNAs as a part of the *P. aeruginosa* Fe-sparing response (17, 18, 51, 52).

**Figure 1.**
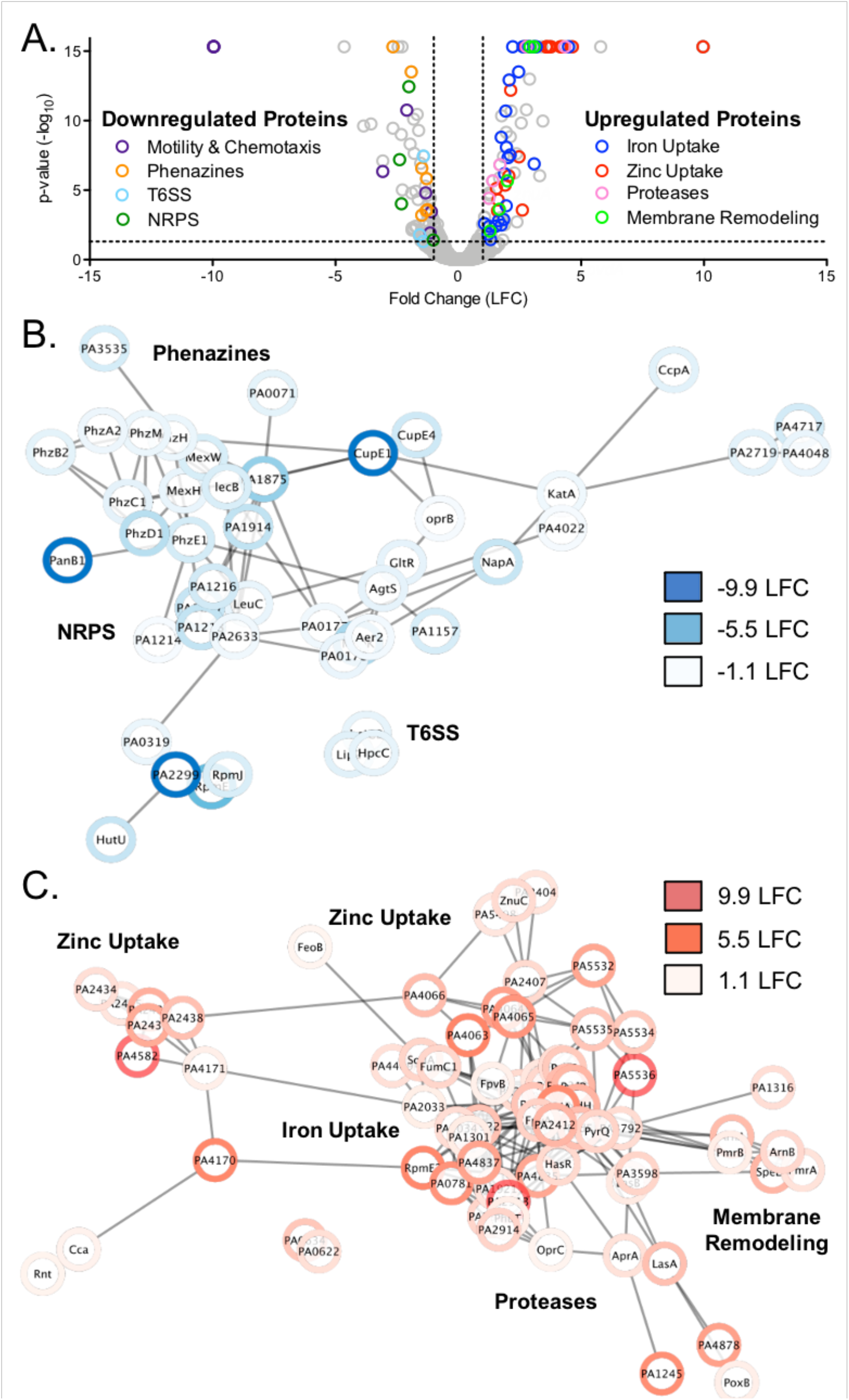
Proteomic analysis of *P. aeruginosa* response to CP. **A.** Volcano plot of PA14 response to CP treatment. Every protein detected in the experiment is represented by an open, gray circle. Circles representing significantly regulated proteins in functional categories of interest are highlighted with the colors indicated in the legend. The thresholds for significance (p<0.05) and expression (1 log_2_ fold change (LFC), equivalent to a 2-fold change) are marked by a black dashed line. Proteomics was performed with an n=5 and significance was determined by ANOVA test corrected for multiple testing by applying a Benjamini-Hochberg procedure (53). **B-C.** Network analysis was performed using the STRING Database (50) on the proteins that were significantly downregulated (**B.**) and upregulated (**C.**) in response to CP treatment. The distance between proteins indicates the strength of the data supporting the interaction with more data correlating to a shorter distance. The network was transferred into Cytoscape (54) and proteomics expression data was integrated using the Omics Visualizer app (55).

Consistent with a recent study (39), CP-treated cells showed a robust Zn-starvation response. This response included increased levels of proteins for Zn(II)-uptake systems, including ZnuACD and proteins for the synthesis and uptake of pseudopaline (CntM and CntO, respectively) (22, 25, 26). CP treatment also led to an increase in several predicted Zn(II)-uptake proteins (PA4063-PA4066, PA2912-PA2914, PA1921, PA1922) that were shown in a previous transcriptomic study to be induced by Zn starvation (25, 26). Also consistent with the previously described Zn-starvation response (23), Zn(II)-containing proteins RpmE and RpmJ were downregulated and Zn(II)-independent proteins RpmE1, DksA2, and PyrC2 were upregulated upon CP treatment (**Fig S2**). Despite the robust Zn-starvation response to CP observed here and in previous work (39), only negligible changes in cell-associated Zn levels were observed for PAO1 and PA14 grown in metal-replete CDM supplemented with 10 μM CP (10, 46).

Although CP has the ability to withhold Mn, levels of known Mn co-factored proteins (HutG, GpmI, UbiD) and predicted Mn-uptake proteins (MntH1, MntH2) in PA14 were not significantly affected by CP treatment. Prior work revealed that the ability of CP to reduce cell-associated Mn in *P. aeruginosa* is medium dependent (10, 46). Whereas CP treatment reduced cell-associated Mn levels when PA14 and PAO1 were grown in a mixture of Tris buffer and tryptic soy both (Tris:TSB) (10), a negligible change in cell-associated Mn was observed when these strains were treated with CP in metal-replete CDM (46). This medium effect may explain the lack of a putative Mn-starvation response in this proteomics analysis. As noted above, the Mn-dependent superoxide dismutase SodM was upregulated in response to CP, possibly resulting from an Fe-starvation response.

In addition to known Fe- and Zn-responsive proteins, proteins for the type VI secretion system (T6SS) (Lip2.1, HsiC2, HcpC), chemotaxis (Aer, Chew2, CheA2, FlgA, CupE2, CupE4), and a non-ribosomal peptide synthetase (NRPS) of unknown function (PA1214, PA1216, PA1217, PA1218) were downregulated upon CP treatment (**Fig 1, Table S1**). CP treatment also led to the upregulation of proteins involved in membrane remodeling and tolerance to CAMPs (PmrA, PmrB, ArnA, ArnB, SpeE2). Curiously, CP treatment caused an increase in several Zn(II)-dependent metalloproteases (ImpA, AprA, LasA, and LasB), contrasting with the Zn-sparing response described above. Altogether, this proteomics analysis indicates that CP exposure induces both Fe- and Zn-starvation responses by *P. aeruginosa* and alters the expression of proteins involved in several virulence processes.

### CP elicits Fe- and Zn-starvation responses that are distinct from single-metal limitation

We next evaluated whether the changes in protein expression described above for CP-treated PA14 resulted from sequestration of an individual metal, multi-metal sequestration, or a metal-independent response to CP using the same quantitative proteomics workflow. We focused these analyses on the limitation of Mn, Fe, and Zn, since the ability of CP to sequester Mn(II), Fe(II) and Zn(II) and induce microbial responses to the limitation of these metal nutrients is established (7, 10, 56). For these experiments, CDM was prepared in the absence of either Fe, Mn, or Zn or lacking all three metals to afford “Fe-depleted”, “Zn-depleted”, “Mn-depleted” and “metal-depleted” CDM, respectively, and protein abundance was compared to that of PA14 grown in metal-replete CDM. Network analysis was used as described above to determine biological connections within the upregulated and downregulated proteomes in response to Fe, Zn, and Mn limitation (**Table S1**).

The Fe-starvation response observed following growth in Fe-depleted CDM was consistent with the well-known regulatory response of *P. aeruginosa* to Fe limitation (17, 51, 52) (**Fig S1**), despite the use of a lower Fe concentration in the metal-replete condition (5 μM instead of 100 μM), a chemically-defined medium [CDM instead of Chelex-treated and dialyzed TSB (DTSB)], a different timepoint (early instead of late stationary phase), and a different strain of *P. aeruginosa* (PA14 versus PAO1). Specifically, Fe limitation resulted in the downregulation of Fe-containing TCA cycle enzymes (SdhBAC, AcnA, PA0794, PA4330), a putative bacterioferritin (PA4880), and Fe-containing oxidative stress response proteins (SodB, KatA); the upregulation of Fe-independent homologs of TCA cycle enzymes (MqoA, FumC1) and a superoxide dismutase (SodM); and the upregulation of pyoverdine- and heme-dependent Fe-uptake systems. When PA14 was grown in Zn-depleted CDM, a Zn-starvation response was observed that was consistent with the transcriptomic response to Zn limitation and investigation of Zn(II)-uptake mutants described in previous investigations (**Fig S2**) (25, 26). Specifically, we observed the downregulation of Zn(II)-containing proteins (RpmE1, RpmJ1) and the upregulation of Zn(II)-independent paralogs of Zn(II)-containing proteins (RpmE2, DksA2, PyrC2, CynT2 and FolE2). CP treatment also led to upregulation of proteins for the biosynthesis and transport of pseudopaline (CmtM, CmtL, CmtO), ZnuABC, the Zn(II)-responsive HmtA heavy-metal-transport system, and several predicted Zn(II)-uptake systems (PA4063-PA4066, PA2912-2914, PA1921). These results demonstrate that the gowth conditions used here and in previous studies to investigate CP (10) allow for characteristic *P. aeruginosa* Fe- and Zn-starvation responses.

When PA14 was grown in Mn-depleted CDM, we found no evidence for a putative Mn-starvation response. The predicted Mn(II) transporters MntH1 and MntH2 were not significantly upregulated in Mn-depleted CDM, and neither known nor predicted Mn-containing proteins were downregulated beyond the established LFC threshold (30). We noted a significant downregulation of SodM, although the fold-change in this protein was below our threshold of 1 LFC (0.8 LFC). Additionally, there was no significant biological connection between the upregulated or downregulated proteins identified by network analysis (**Table S2**). Because no robust Mn-starvation response was observed either upon Mn-limitation or CP treatment, we focused the remainder of our metal-depletion studies on Fe and Zn.

We next identified overlaps in the proteomic responses to CP treatment, Fe limitation, and Zn limitation. We observed ~35% overlap between proteins upregulated by CP treatment and Fe limitation, including proteins involved in the uptake of Fe(III)-pyoverdine and heme (**Fig 2B, Table S3**). Proteins that were observed to be upregulated by Fe limitation but not CP treatment included several Fe(III)-uptake proteins that were either not detected in the CP treatment experiment (8 proteins) or not significantly upregulated beyond the 1-LFC threshold (PhuR); separate studies are underway to determine whether the undetected proteins are also altered by CP treatment. Approximately 20% of the proteins downregulated in response to Fe limitation were also downregulated in response to CP (**Fig. 2, Table S3**). These proteins included phenazine biosynthesis and secretion proteins and the NRPS operon noted above. Further analysis of the 99 downregulated proteins that appeared specific to Fe limitation revealed that 9 of these proteins were significantly (p<0.05) downregulated upon CP treatment, but the change did not meet the 1-LFC threshold, possibly indicating a weaker Fe-starvation response to CP than induced by Fe-depleted CDM (**Fig S1**). Notably, several PrrF regulon proteins, including TCA cycle enzymes (SdhABCD, AcnB), NADH dehydrogenase (NuoABCEFGHI), and the putative bacterioferritin (PA4880) were not significantly changed in the presence of CP, indicating that CP treatment elicits an incomplete Fe-sparing response. Previous studies using an AntR transcriptional and translational reporter strain showed that AntR, which is downregulated by PrrF during growth in DTSB (57), is also downregulated in response to CP treatment during growth in Tris:TSB (10). However, AntR and its regulatory targets AntABC were not detected in this proteomics experiment. To determine whether the Fe-starvation response to CP during growth in metal-replete CDM included a decrease in AntR, we used the same reporter strain and found that *antR* expression is similarly repressed by CP treatment during growth in CDM (**Fig S2**), indicating that PrrF functions under these conditions. Combined, these data indicate that that CP elicits only a portion of the Fe-starvation response that is observed upon Fe limitation in Fe-depleted CDM.

**Figure 2.**
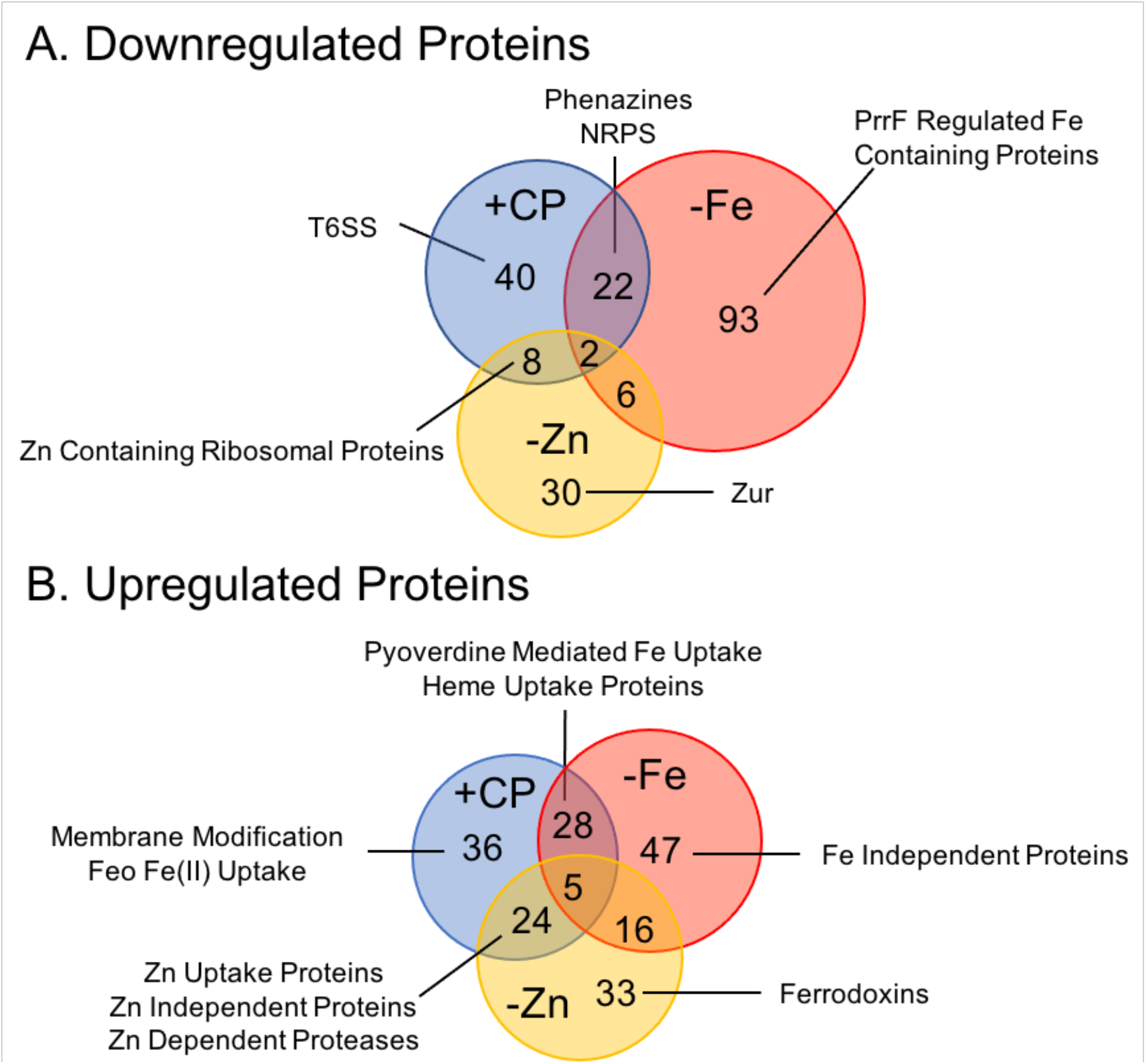
Comparison of the proteome in response to Fe and Zn limitation and CP treatment. Venn diagrams of significantly (p<0.05) downregulated (>1 LFC) (**A.**) and upregulated (>1 LFC) (**B.**) proteins in response to CP treatment (+CP), Fe limitation (−Fe), and Zn limitation (−Zn). Proteomics was performed with n=5; LFC = log_2_ fold change.

The shared response to both Zn limitation and CP treatment was consistent with the known Zn-starvation response and included the downregulation of Zn(II)-containing ribosomal proteins (RpmJ and RpmE) (**Fig 2A, Fig S3**) and the upregulation of Znu transport proteins (ZnuA, ZnuC), pseudopaline biosynthesis and transport (CntM, CntO) proteins, the Zn(II)-independent ribosomal protein RpmE1, and the Zn(II)-independent paralog to DksA, DksA2 (**Fig 2B, Fig S3**). Notably, several Zn(II)-dependent proteases (LasA, LasB, ImpA, AprA) were upregulated by both CP and Zn(II) limitation, seemingly contrary to the observed Zn-starvation response and a recent report on decreased protease activity in response to CP (39) (**Fig 2B**). The upregulated proteins that were specific to the Zn-limitation proteome included Zn(II)-independent proteins (CynT2, FolE2) that were not significantly changed in response to CP. Moreover, two ferredoxins (Fdx2, FdxA) were upregulated by Zn limitation and were not significantly affected by CP treatment (**Table S3**). Notably, the Zn(II)-responsive transcriptional regulator Zur was downregulated only in response to Zn limitation. Previous work demonstrated *zur* is co-transcribed with *znuC* and *znuB*, and expression of all three genes is induced by Zn limitation (**Fig S4A**) (22). However, in the current work, Zur was downregulated by Zn limitation and unchanged in response to CP, whereas ZnuC and ZnuB were upregulated in both the Zn-depeleted and CP-treatment conditions (**Fig S4B**). We performed real-time PCR (RT-PCR) analysis to investigate this finding further and observed that expression of both the *znuA* and *zur* mRNAs were induced by CP treatment and by Zn limitation (**Fig S4C-D**), suggesting that Zur is post-transcriptionally downregulated upon Zn limitation and CP treatment. Together, these observations suggest that the response of *P. aeruginosa* to Zn starvation is more complex than currently appreciated; this notion warrants further investigation.

The comparisons between the responses of PA14 to CP, Fe limitation, and Zn limitation also identified responses that were unique to CP treatment. Several T6SS proteins encoded by the HSI-II T6SS locus (**Fig 2A**, **Fig 3A**) were downregulated only in response to CP. This result was surprising given that our recent proteomic studies showed that Fe starvation upregulates the expression of the same T6SS proteins (47). However, the mRNAs encoding three of the HSI-II T6SS proteins – *lip2, clipV2*, and *hsiB2* – were not significantly changed in a subsequent RT-PCR experiment (**Fig 3B-D)**, contrasting with the recent Fe-regulation study (47) and the current proteomics results (**Fig 3A**). The difference in gene expression between this work and the previous study may result from differences in experimental conditions and post-transcriptional effects may be responsible for the distinct effects of CP on protein expression. Also notable was the Feo Fe(II)-import system, which was upregulated only in response to CP but not in response to Fe limitation (**Fig 2A, Fig S1**). Additionally, proteins involved in membrane modification and CAMP resistance (PmrA, PmrB, ArnA, ArnB, SpeE2) were upregulated only in response to CP (**Fig 2B, Table S3**). The Feo Fe(II)-transport system has been shown to be regulated by PmrAB (58), possibly explaining the increase in its expression in response to CP. Together, these data indicate that CP treatment elicits a physiological response that overlaps with but is distinct from the *P. aeruginosa* Fe- and Zn-starvation responses.

**Figure 3.**
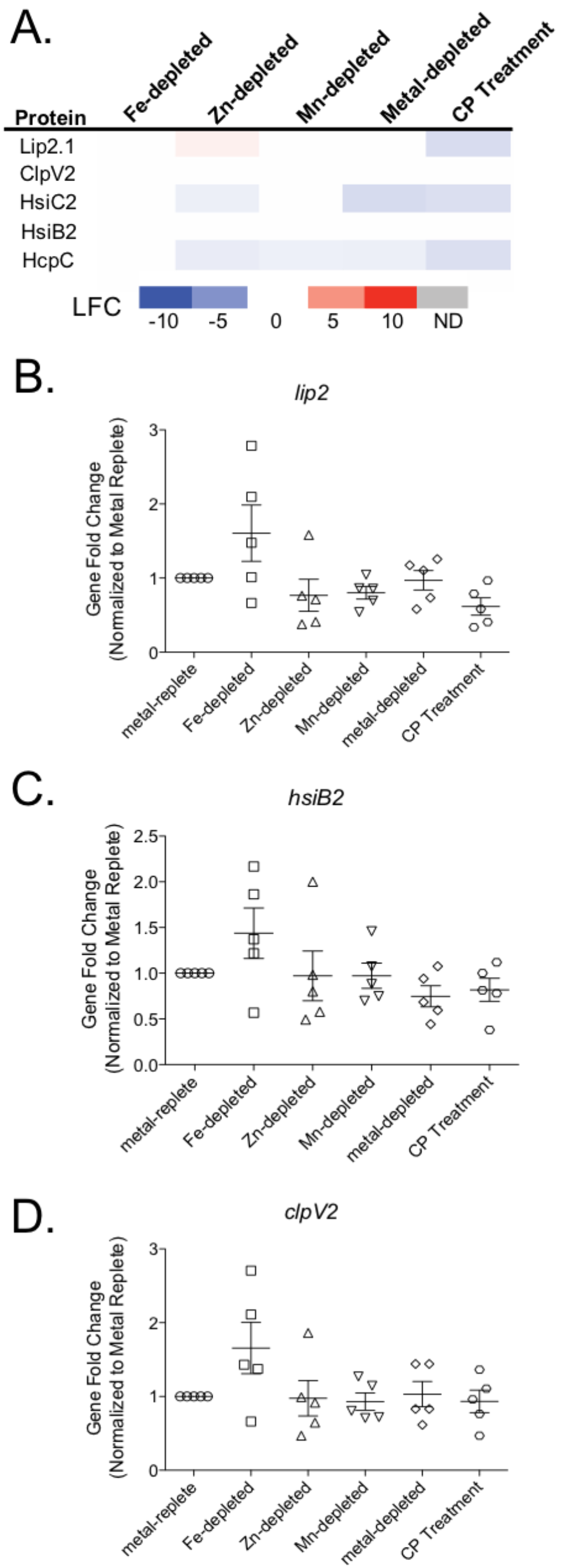
T6SS proteins are downregulated in response to CP. **A.** Heatmap of protein expression of *P. aeruginosa* PA14 grown in Fe-depleted, Zn-depleted, Mn-depleted, metal-depleted CDM, and in metal replete CDM in the presence of CP compared to expression in metal-replete CDM. The log_2_ fold change (LFC) is shown for all significantly (p<0.05) changed proteins. Gene expression under the same conditions was measured for *lip2* (**B.**), *hsiB2* (**C.**), and *clpV2* (**D.**) using RT-PCR. No significance was detected in any of the comparisons. Significance was determined by one-way ANOVA with Dunnet’s multiple comparisons test. n=5.

### CP treatment and Zn limitation reduce protease activity despite increases in protease levels

The observed increase in the levels of secreted proteases (LasB, LasA, AprA, ImpA) in response to Zn limitation and CP treatment (**Fig. 4A**) was surprising for several reasons. First, these proteases are secreted proteins, and our workflow was designed to analyze the cell-associated proteomes of *P. aeruginosa*. Second, a previous study demonstrated decreased protease activity under Zn-limiting conditions (26), and more recent work showed a similar decrease in protease activity upon chelation of Zn(II) by CP (39). To investigate this finding further, we quantified the secreted protease activity against azocasein of PA14 grown in metal-replete, Fe-depleted, Zn-depleted, Mn-depleted, and metal-depleted CDM, or in the presence of CP. In contrast to what was observed with cell-associated protease levels and in agreement with previous studies (26, 39), protease activity in the culture supernatants decreased upon both Zn limitation and CP treatment (**Fig 4B**). A previous study with PAO1 reported the restoration of protease activity in the Δ*znuA* and Δ*cntO* Zn(II)-uptake mutants with the addition of Zn(II) to the assay buffer but not in Δ*znuA*Δ*cntO* (26). Similar to the Δ*znuA*Δ*cntO* mutant, we were unable to recover protease activity to Zn-depleted or CP-treated culture supernatants by supplementing the assay buffer with Zn(II) (**Fig 4C**). The Δ*znuA*Δ*cntO* double mutant had a marked growth defect during growth in minimal media and significantly lower cell-associated Zn when compared to wild type and the single mutants, indicating it was more Zn starved than the single mutants (26). Our current results indicate that Zn-depleted and CP-treated cultures were similarly too Zn-starved to recover protease activity with the addition of Zn(II) to the assay buffer.

**Figure 4.**
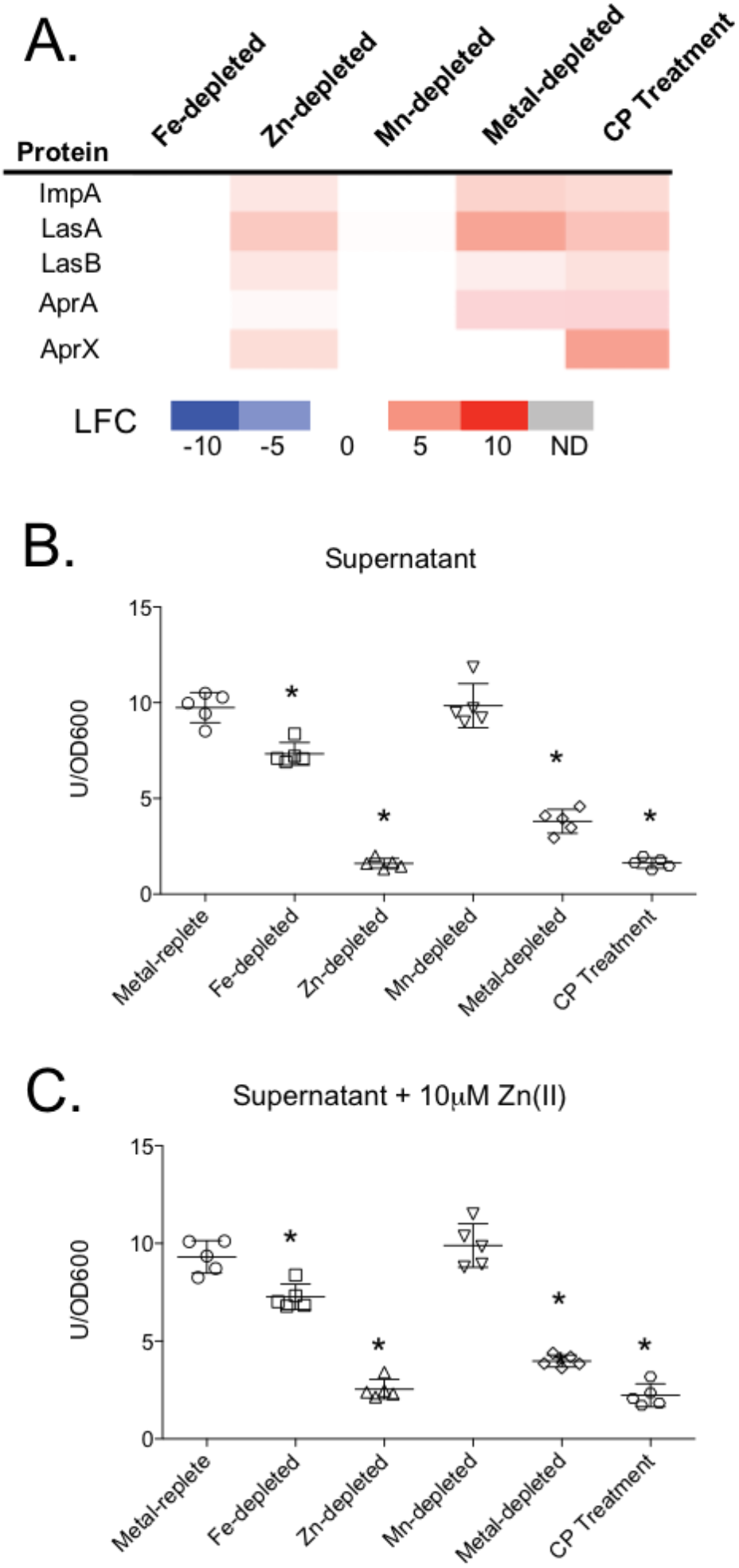
Secreted protease activity decreased in response to Zn limitation despite an increase in cell associated protein abundance. Heatmap (**A.**) of secreted proteases shown for all experimental conditions compared to the metal replete control. The log_2_ fold change (LFC) of significantly (p<0.05) changed proteins is shown for selected proteins. The secreted protease activity was measured in the absence (**B.**) and presence (**C.**) of 10 μM Zn(II) in the assay buffer to determine if activity could be restored. Activity was determined by azocasin assay and normalized to OD_600_ after 8 hours of growth. Significance was determined by one-way ANOVA with Dunnet’s multiple comparisons test. n=5, *p<0.05.

While these results were generally consistent with previous studies, they did not explain the increase in abundance of the cell-associated proteases. We considered that the expression of secretory systems responsible for export each of these proteases may be downregulated. LasA, LasB, and ImpA are secreted by the Xcp and Hxc type II secretion systems (T2SS) (59, 60), whereas the alkaline protease AprA is secreted by a type I secretion system (T1SS) encoded by the *aprDEF* operon upstream the *aprA* gene (61). Amongst the proteins for each of these systems, our proteomics data showed that only two proteins – XcpX and AprF – were downregulated upon CP treatment. Upon Zn limitation, XcpX was not significantly changed and AprF was downregulated below our threshold of 1 LFC (**Fig S5A**). In contrast, HxcV was strongly repressed when PA14 was cultured in Zn-depleted medium. This protein was not detected in the CP-treated samples; thus, we cannot exclude the possibility that CP similarly repressed its expression. Further gene expression analysis showed that expression of *xcpT* did not change upon CP treatment or Zn limitation, whereas *xcpP* expression increased a small but statistically significant amount in response to both CP treatment and Fe limitation (**Fig S5B, Fig S5C**). Similarly, expression of genes within the *xcp* operons were not significantly changed under any condition tested (**Fig S5D-G)**. Overall, these data suggest that decreased expression of proteins that secrete Zn(II)-dependent proteases is unlikely to contribute to the decreased protease activity in culture supernatants despite increased levels of these proteins within cells.

### CP treatment increases levels of membrane-remodeling proteins

As mentioned above, CP treatment resulted in increased expression of the two-component regulator PmrAB and its regulatory targets ArnA and ArnB, which synthesize aminoarabinose, and the spermidine synthetase SpeE2 (**Fig 5A**) (62, 63). The addition of spermidine to lipopolysaccharide (LPS) and aminoarabinose to lipid A promotes resistance to CAMPs, including colistin (polymyxin E) and polymyxin B (62, 64, 65). Initially, it appeared that the induction of membrane-remodeling machinery did not overlap with the Fe- or Zn-starvation responses and therefore may be part of a metal-independent response. Further analysis, however, showed that Zn limitation caused a small but significant increase in SpeE2 that fell below our initial LFC threshold of 1 (0.6 LFC). Moreover, SpeE2, ArnB and PmrA were significantly upregulated in metal-depleted CDM, suggesting that this CP-induced response resulted from multi-metal sequestration. To investigate whether CP treatment afforded CAMP resistance in *P. aeruginosa*, we evaluated how CP pre-treatment of PA14 affected the minimal bactericidal concentration (MBC) of polymyxin B. Because previous studies demonstrated increased polymyxin B resistance after growth in low-Mg(II) conditions that results from aminoarabinose modification of lipid A (58), PA14 was also pre-cultured in metal-replete CDM supplemented with 20 μM Mg(II) (low-Mg CDM), instead of the standard 2 mM Mg(II), as a positive control. Although we noted some variation in MBCs between three biological replicates, bacteria pre-cultured in metal-replete CDM were consistently less resistant to polymyxin B than bacteria pre-cultured in metal replete, low-Mg CDM (**Fig 5B, Table S4**). PA14 pre-cultured in either metal-depleted CDM or pre-treated with CP exhibited MBC ranges (64-128 mg/L and 32-128 mg/L, respectively) that overlapped with that of the cultures grown in low-Mg CDM (32-64 mg/L). Statistical significance (p<0.05) was identified between the metal-depleted and metal-replete samples by Student’s *t* test. However, significance was not found between the experimental conditions (pre-cultured in metal-depleted CDM, low-Mg CDM, or CP treatment) and the control condition (metal-replete CDM) by one-way ANOVA with Dunnets post-test. More in-depth investigations of this finding are currently underway. Together, these data suggest that metal withholding and CP increases the resistance of PA14 to polymyxin B.

**Figure 5.**
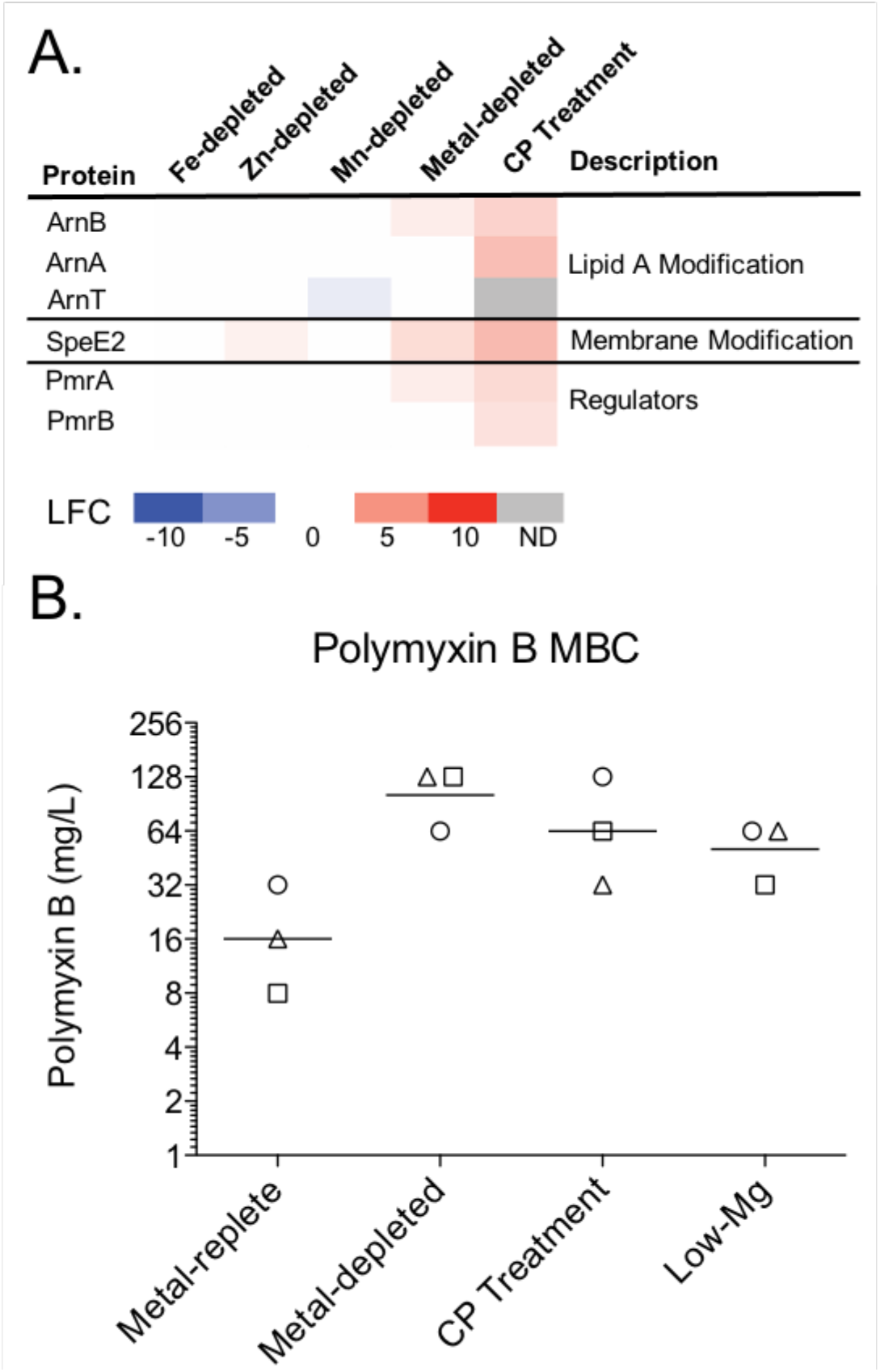
Polymyxin B resistance increases after exposure to CP. **A.** Heatmap of membrane-remodeling proteins under all metal-depletion conditions. Protein expression is compared to the metal-replete condition. The log_2_-fold change (LFC) of each significantly (p<0.05) changed protein is shown. **B.** The minimal bactericidal concentration (MBC) of polymyxin B was determined for PA14 after pre-culture in metal-replete, metal-depleted, and low-Mg(II) CDM, and after CP treatment. The geometric mean of the three biological replicates for each condition is denoted with a line. The biological replicates are represented with circles (replicate 1), squares (replicate 2), and triangles (replicate 3).

## Discussion

In the current work, we systematically evaluated the contributions of Fe-, Zn-, and Mn-limitation to the proteomic response of *P. aeruginosa* to CP in a chemically-defined medium. We previously demonstrated an Fe-starvation response to CP (10) but reasoned that a broader physiological response was necessary to maintain metal homeostasis in response to CP multi-metal withholding. Our results demonstrate that the response of *P. aeruginosa* to CP is not simply an additive response to Fe and Zn withholding. Moreover, we identified CP-dependent induction of membrane-modification proteins that appear to afford polymyxin B resistance. Since this response was not observed upon depletion of any single metal, we hypothesize it is dependent upon a general depletion of transition metals by CP. Consequently, this work demonstrates that multi-metal sequestration elicits distinct effects as compared to individual metal depletion on the physiology and virulence capacity of a human pathogen.

A goal of our study was to determine how sequestration of Fe, Zn, and Mn contribute to the response of *P. aeruginosa* to CP. Thus, in our analyses, specific responses to single-metal limitation were compared to the response to CP multi-metal withholding. This analysis revealed that the CP-induced Fe-starvation response was markedly incomplete compared to the Fe-starvation response observed in Fe-depleted medium. For example, CP induced upregulation of Fe-acquisition proteins, but many Fe-containing metabolic proteins, which are normally downregulated by the PrrF sRNAs upon Fe limitation (52), were not significantly affected by CP treatment. Consistent with our findings, a previous transcriptomic study of the response of *P. aeruginosa* to CP showed decreased expression of phenazine biosynthesis genes and increased expression of pyoverdine biosynthesis genes, yet no change in the expression of PrrF-responsive TCA cycle genes (38). This earlier study attributed many of these changes to Zn limitation, since the Fe(II)-withholding ability of CP was not yet accepted, leading to the proposal that a subset of Fe-regulated genes could also be responsive to other metals (38). Nevertheless, our previous proteomics study of Fe-limited PAO1 demonstrated that decreased expression of phenazine biosynthesis proteins and increased expression of pyoverdine biosynthesis proteins occurred in a strictly Fe-dependent manner, suggesting that CP induces some but not all of the prototypical Fe-starvation response in *P. aeruginosa* (10). The current study further supports that decreased expression of phenazine biosynthesis proteins and increased expression of pyoverdine biosynthesis proteins result from Fe limitation. The origin(s) of the incomplete Fe-starvation response to CP is unclear. We currently speculate that it results from a tiered response to different levels of Fe limitation similar to what was found for the *Bacillus subtilis* Zn-starvation response (66). For example, the Zn-starvation response that is elicted by CP treatment may mask the activation of specific Fe-starvation pathways, resulting in a distinct Fe-starvation response upon CP treatment as compared to Fe limitation. More work is needed to understand the observed differences in Fe-starvation responses between these two conditions.

We also observed a robust Zn-starvation response to CP, evidenced by the upregulation of multiple Zn(II)-uptake systems by PA14. In contrast to the incomplete CP-induced Fe-starvation response described above, the Zn-starvation response elicited by CP was comparable to that caused by Zn limitation, suggesting a robust response to Zn(II) withholding by CP. Notably, the Zn-starvation response is induced in the absence of a decrease in cell-associated Zn during CP treatment (10, 46). This observation is consistent with previous reports that CP treatment results in metal-starvation responses without changes to the corresponding cell-associated metal concentrations (67, 68). For instance, CP was found to exert antimicrobial activity and induce a Zn-starvation response in *Candida albicans* without decreasing cell-associated Zn (67). In *A. baumannii* the Cu uptake protein OprC was downregulated and a Cu storage protein was upregulated, indicating a Cu-starvation response to CP, without a change in cell-associated Cu (68). One reason for these differences is that the measurements provide the total cellular metal content, which includes both labile metal pools that can be sensed by the cell as well as kinetically inaccessible (e.g. stored) metal pools. Together, these data indicate that a decrease in cell-associated metal content is not always a reliable indicator of how pathogens respond to multi-metal withholding.

In addition to the Zn-starvation response, CP treatment and Zn limitation both resulted in an unexpected increase in Zn(II)-dependent proteases. Further analyses showed that CP treatment and Zn limitation decreased protease activity in culture supernatants, aligning with previous studies (24, 26, 39, 69). Because of the intracellular increase in abundance of these secreted proteins, we initially hypothesized that a decrease in protease secretion may contribute to the observed decrease in protease activity. However, we observed little change in protein abundance or in transcript levels for the Xcp and Hxc T2SS systems that export these proteases, though it is possible that protease secretion is reduced by an unknown mechanism. A previous study demonstrated that CP chelated Zn(II) from proteases when supernatant was treated with CP (39). Further work will be needed to determine the mechanism of decreased protease during Zn limitation and to determine the mechanism of decreased proteased activity during CP treatment of cultures.

Recently, CP was shown to induce a multi-metal starvation response in *A. baumannii* (68). In addition to the induction of siderophores and other Fe-uptake systems, the authors proposed a dual-regulatory system that responds to Zn withholding and flavin mononucleotide (FMN) concentration to induce a Zn(II)-independent enzyme that maintains flavin biosynthesis during Zn limitation. Flavin biosynthesis is necessary for the production of flavodoxins and FMNs, which can act as Fe-independent substitutes for ferrodoxins and cofactor Fe-independent homologs, respectively. Our own recent work identified potential cross-talk between the Fe and Zn homeostasis systems of *P. aeruginosa* (52), but this cross-talk was not evident in the current study, possibly due to the 20-fold lower Fe concentration used for the proteomics experiments. Thus, it remains unclear how much the interactions of distinct metal-regulatory systems contribute to the response of *P. aeruginosa* to CP.

We also identified what appears to be a multi-metal limitation response upon CP treatment with the induction of the PmrAB two-component regulatory system and its targets ArnA, ArnB, and SpeE2 (**Fig 2, Fig 5**). PmrAB activation of the *arn* operon results in the addition of aminoarabinose modifications to lipid A, resulting in increased resistance to CAMPs such as colistin and polymyxin B. The PmrAB regulon is known to be induced by Mg(II) and Ca(II) limitation (62, 70). Here we provide evidence that this operon is more generally responsive to low concentrations of transition metals, including Fe, Mn, and Zn limitation caused by CP. A previous study demonstrated that extracellular DNA binds cations and induces resistance to CAMPs through the activity of PmrAB regulated proteins (71), further supporting the conclusion that a more generalized low-cation environment can initiate the PmrAB pathway. We further provide evidence that both CP treatment and transition-metal depletion results in increased resistance to polymyxin B. While we did not address the mechanism in this study, the observed upregulation of ArnA and ArnB suggests that the increased resistance occurs through the addition of aminoarabinose to lipid A (62, 64) This modification is highly relevant in the clinical setting as aminoarabinose modification of lipid A is commonly observed in polymyxin-resistant *P. aeruginosa* isolates (72, 73). Nebulized colistin is commonly used to reduce *P. aeruginosa* burdens in the lungs of CF patients (74–76). Our results suggest that CP released during the inflammatory response in the CF lung may promote *P. aeruginosa* to resist this treatment. CP also decreased activity of lipid A biosynthesis proteins in *Helicobacter pylori*, resulting in altered lipid A structure which led to increased biofilm production and reduced growth inhibitory activity of CP (77). It is possible that changes to lipid A are a widespread response of Gram-negative bacteria to CP.

In closing, the results from this proteomics study provide new insights into host-pathogen interactions by systematically evaluating how multi-metal withholding by CP impacts *P. aeruginosa* physiology. This work demonstrates the response of *P. aeruginosa* to metal withholding by CP is not the additive response of individual metal-starvation responses. Moreover, metal-starvation responses are linked to virulence regulatory networks in many pathogens, and the current work highlights that response of *P. aeruginosa* to multi-metal limitation elicits several changes in virulence-related processes. Thus, this investigation broadens our understanding of how nutritional immunity impacts the pathogenic potential of an important opportunistic bacterial pathogen.

## Materials and Methods

### Growth media and conditions

*P. aeruginosa* strain PA14 was used for all experiments. Throughout the paper we have used the locus tags for PAO1 instead of PA14 when gene names are not available for ease of reading and comparisions to the literature. PA14 was grown as previously described in chemically-defined media (CDM). CDM was made as previously described (10, 40) with some modifications (**Table S4**). Briefly, the medium was made without added Mg(II), Ca(II), and was then aliquoted and stored at −80°C. Prior to an experiment, the medium was thawed and supplemented with 0.1 μM NiCl_2_, 0.1 μM CuCl_2_, 5 μM FeSO_4_, 6 μM ZnCl_2_, 0.3 μM MnCl_2_, 2 mM MgSO_4_, and 2 mM CaCl_2_ to afford metal-repleteCDM. Calprotectin (CP, 10 μM final concentration) was added to CDM where indicated. Metal-depleted CDM was made by supplementing only with NiCl_2_, CuCl_2_, MgSO_4_, and CaCl_2_. Fe-depleted, Zn-depleted, and Mn-depleted CDM were made by not adding Fe, Zn, or Mn, respectively, to CDM. Low-Mg CDM was made by adding 20 μM MgSO_4_ instead of 2 mM MgSO_4_ to metal-replete CDM. Cultures for all experiments were inoculated to and OD_600_ of 0.05 and grown for 8 hours shaking at 250rpm at 37°C.

### Calprotectin purification

CP was purified as previously described (78). Protein aliquots were stored in 20 mM HEPES, 100 mM NaCl, 5 mM DTT, pH 8.0 at −80°C. Aliquots were thawed only once before use and buffer exchanged three times into 20 mM Tris, 100 mM NaCl, pH 7.5 using pre-sterilized 10K MWCO spin concentrators (Amicon). Protein concentrations were determined by A_280_ using the calculated extinction coefficient of the CP heterodimer (ε_280_=18540 M^−1^cm^−1^) obtained from the online ExPASy ProtParam tool.

### Quantitative label-free proteomics

Two independent proteomics experiments were performed each with five biological replicates. The media for both experiments was prepared as described above as a single batch, inoculated with the same five overnight cultures, and samples were collected for both experiments at the same time to limit variability. For the first experiment, PA14 was grown in metal-replete CDM with and without 10 μM CP. For the second experiment, PA14 was grown in metal-replete, Fe-depleted, Mn-depleted, Zn-depleted, and metal-depleted CDM. Quantitative label-free proteomics was performed using a Waters nanoACQUITY UPLC coupled to a Thermo Orbitrap Fusion Lumos Tribrid mass spectrometer similar as previously described with modification (47–49). A full description of the mass spectrometry-based experimental methodology is provided in the Supplementary Information. Gene function and pathway analysis was conducted using information from the *Pseudomonas* genome database (79) and the *Pseudomonas* metabolome database (80), and the STRING database (50).

### Network analysis

Network analysis was performed using the STRING Database version 10.5 (50). Corresponding PAO1 accession numbers were used as the database is limited to the PAO1 strain. The network was downloaded from STRING and further analyzed using Cytoscape (54). The Omics Viewer app was used within Cytoscape to incorporate proteomics expression data into networks (55).

### AntR reporter assay

PA14/P*antR*-‘*lacZ-*SD was generated previously (10). Cultures were grown in CDM with or without 10 μM CP as described above. β-Galatosidase activity was measured as previously described (81). Briefly, bacterial growth was measured by OD_600_, cells were harvested by centrifugation and resuspeneded in potassium phosphate buffer (50 mM, pH 7.0). Cells were diluted 1:10 in Z-buffer (60 mM Na_2_HPO_4_, 35 mM NaH_2_PO_4_, 1 mM KCl, 100 mM MgSO_4_, 50 mM β-mercaptoethanol) and lysed using chloroform and 0.1% SDS. The enzyme reaction was started using *o*-nitrophenyl-β-D-galactopyranoside (ONPG, 4 mg/mL) dissolved in potassium phosphate buffer and stopped using sodium carbonate (1 M). The quenched reaction was centrifuged to remove cell debris and the supernatant absorbance was read at A_420_. β-galatosidase activity was calculated into Miller Units using the following equation: MU = (1000 x A_420_) / (time [mins] x culture volume [mL] x OD_600_).

### Real-time PCR

Five cultures of PA14 were grown as described above in CDM, Fe-depleted, Mn-depleted, Zn-depleted, and metal-depleted CDM, and in CDM in the presence of CP. Real-time PCR (RT-PCR) was performed as previously described (21). Briefly, cell pellets were stored in RNA*Later* at −80°C. RNA was extracted using a Qiagen RNeasy kit, cDNA was synthesized, and RT-PCR was performed using Taqman reagents (Roche) using a StepOnePlus (ThermoFisher). Relative expression was determined using the ΔΔCt method. Expression was normalized to the 16S ribosomal gene. Primers and probes used are listed in **Table S5**.

### Azocasein protease activity assay

Five biological replicates of PA14 were grown as described above. After 8 hours of growth, cultures were centrifuged at 16,000 x g for 10 minutes. The supernatants were sterile filtered using 0.2-μm-pore-size filters (Costar) and stored at 4°C overnight. Total protease activity was quantified as previously described (82). Briefly, 20 μL of supernatant was added to 0.5 mL of the 0.3% azocasein solution (50 mM Tris-HCl (pH 7.2) 0.5 mM CaCl_2_ buffer and supplemented with or without 10 μM ZnCl_2_) and incubated for 30 minutes at 37°C. The reaction was stopped with 0.5 mL 10% tricholoracetic acid. The quenched reaction mixture was centrifuged at 16,000 x g for 20 minutes and the supernatant absorbance was read at 400 nm using a Biotek Synergy HT plate reader. One unit of enzyme activity changes the A_400_ by 0.01. The enzyme activity was normalized to OD_600_.

### Polymyxin killing assay

Three biological replicates of PA14 were grown as described above in CDM, metal-depleted, and low-Mg CDM, and in CDM in the presence of 10 μM CP. After 8 hours of growth, the cultures were centrifuged, and the cells were resuspended in PBS. The OD_600_ was measured and PBS was inoculated to an OD_600_ of 0.05, which was subsequently diluted 1:10 into CDM with no CaCl_2_ supplementation. The 1:10 dilution was used to inoculate 1×10^5^ CFU into 2 mL CDM with no CaCl_2_ supplementation containing 0, 1, 2, 4, 8, 16, 32, or 64 mg/L polymyxin B. Serial dilutions of the inoculum were performed and plated on Pseudomonas Isolation Agar (PIA) and incubated overnight to ensure inoculum was at 1×10^5^ CFU/mL. Cultures were incubated for 18 hours at 37°C. After growth, 10 μL of culture was spotted onto PIA, and the plates were incubated overnight at 37°C.

## Acknowledgements

This work was supported by the NIH (R01 GM126376 to E.M.N. and A.G.O.; T32 AI095190 to C.E.N.) and the University of Maryland School of Pharmacy Mass Spectrometry (SOP1841-IQB2014 to MAK). E.M.Z. is a recipient of a NSF Graduate Research Fellowship and a Whitaker Health Science Fund Fellowship.

